# Lipid binding by N-terminal motif mediates plasma membrane localization of *Bordetella* effector protein BteA

**DOI:** 10.1101/2020.11.17.386359

**Authors:** Ivana Malcova, Ladislav Bumba, Filip Uljanic, Darya Kuzmenko, Jana Nedomova, Jana Kamanova

**Author notes:** Corresponding author (JK). These authors contributed equally to this work (Ivana Malcova, Ladislav Bumba).

## Abstract

The classical *Bordetella* species, *B. pertussis* and *B. bronchiseptica*, employ a type III secretion system (T3SS) to inject a 69-kDa BteA effector into the host cells. Upon injection, BteA localizes to the cytosolic leaflet of lipid rafts via its N-terminal lipid raft targeting (LRT) domain and induces cell death. The plasma membrane targeting and cytotoxicity mechanisms of BteA are poorly understood. Using protein-lipid overlay assay and surface plasmon resonance, we showed here that the recombinant LRT domain, which adopts a four-helix bundle topology of membrane localization domains, specifically binds negatively charged membrane phospholipids. The binding affinity for phosphatidylinositol 4,5-bisphosphate (PIP2)-containing liposomes with Kd ~450 nM was higher than for those enriched in phosphatidylserine (Kd ~1.2 μM) while both phospholipids were required for plasma membrane targeting in yeast cells. The membrane association of LRT further depended on its electrostatic and hydrophobic interactions and involved a loop L1-located leucine residue. Importantly, charge-reversal substitutions within the L1 region disrupted plasma membrane localization of BteA effector without hampering its cytotoxic activity during *B. bronchiseptica* infection of HeLa cells. The LRT-mediated targeting of BteA to the cytosolic leaflet of the plasma membrane of host cells is, hence, dispensable for the effector cytotoxicity.

**Author summary:** The respiratory pathogens of humans and other animals, *Bordetella pertussis* and *Bordetella bronchiseptica*, produce a type III secretion system effector protein BteA. This effector consists of two functional domains, an N-terminal lipid raft targeting (LRT) domain, and a cytotoxic C-terminal domain, which induces non-apoptotic and caspase-1-independent host cell death. We found here that the LRT domain of BteA associates with plasma membrane by binding to negatively charged phospholipids. We further discovered that the mechanism of membrane association by LRT is reminiscent of the one used by three other diverse families of toxins: clostridial glucosyltransferase toxins, multifunctional-autoprocessing RTX toxins (MARTX), and *Pasteurella multocida-like* toxins. Intriguingly, we also report that plasma membrane targeting by the LRT domain does not contribute to cytotoxic activity of BteA during *B. bronchiseptica* infection. Overall, our work elucidated the mechanism of plasma membrane association by LRT, and further provided the basis for future research on cellular activities of BteA and the mechanism of BteA-induced cell death.

## Introduction

Bacterial toxins and effectors localize to specific compartments within the host cell environment to access their intracellular targets and enhance the signaling specificity and efficacy. Various membrane-targeting strategies have evolved to trap the catalytic domain or adaptor domain at the proximity of the membrane-anchored substrates [1]. Some of the proteins insert directly into the membrane as integral membrane proteins through transmembrane domains, such as the *Salmonella* type III secretion system (T3SS) effector SteD [2]. Others undergo covalent lipid modification, such as *Salmonella* T3SS effector proteins SspH2 and SseI that are S-palmitoylated on a conserved cysteine residue within their N-terminal domains by a specific subset of host-cell palmitoyltransferases [3]. Nevertheless, a significant part of bacterial toxins and effectors possess a dedicated membrane localization domain (MLD) that binds phospholipids. For example, recruitment of the *Legionella pneumophila* effector DrrA to the *Legionella*-containing vacuole, where it AMPylates Rab1, is mediated by a phosphatidylinositol 4-phosphate (PI(4)P)-binding domain. This domain is characterized by a deep electropositive binding pocket and surrounding membrane-penetrating leucine residues [4, 5]. Another domain termed 4HBM for four-helix bundle membrane localization domain is shared by multiple bacterial toxins, including *Pasteurella multocida* mitogenic toxin (PMT), multifunctional-autoprocessing RTX toxins (MARTX), and large clostridial glucosyltransferase toxins exemplified by *Clostridium difficile* toxin A (TcdA) and toxin B (TcdB), and *C. sordellii* lethal toxin (TcsL) [1, 6-8]. The phospholipid-binding site of 4HBM is located at the apex of the structure, the so-called bundle tip, which is formed by two protruding loops, loop 1 (L1) connecting helices 1 and 2, and loop 3 (L3) in between helices 3 and 4. Indeed, the positively charged and hydrophobic residues within L1 and L3 are key for lipid-binding and membrane localization of 4HBM as revealed by their mutagenesis [8, 9]. Furthermore, the proper localization of PMT and TcsL toxins also proved to be critical for both toxin activities and TcsL cytotoxicity [6, 10].

The cytotoxic effector BteA is injected into the host cells by a type III secretion system (T3SS) of classical *Bordetella* species, *B. pertussis* and *B. bronchiseptica* [11]. These bacteria colonize the ciliated epithelia of the respiratory tract of diverse mammals and cause respiratory illness with differing symptoms, duration, and severity. The strictly human-adapted *B. pertussis* is the primary causative agent of pertussis or whooping cough, a contagious respiratory illness of humans that remains one of the least controlled vaccine-preventable infectious diseases [12]. The *B. bronchiseptica* species, on the other hand, infects a broad range of mammals and causes infections ranging from lethal pneumonia to asymptomatic and chronic respiratory carriage [12, 13]. The activity of T3SS of *B. bronchiseptica* is required for persistent colonization of the lower respiratory tract of rats, mice, and pigs, presumably due to the actions of BteA effector [14-16]. In tissue culture, however, BteA actions account for rapid cell death that is non-apoptotic and caspase-1-independent [17]. Remarkably, compared to the high BteA-mediated cytotoxicity of *B. bronchiseptica*, the cytotoxicity of BteA of *B. pertussis* is strongly attenuated due to the insertion of an extra alanine at position 503, which may represent an evolutionary adaptation of *B. pertussis* [18]. Nevertheless, the role of BteA and T3SS activity in the pathophysiology of human pertussis remains to be established.

The 69-kDa effector protein BteA exhibits modular architecture, consisting of two functional domains, an N-terminal localization/lipid raft targeting domain of ~ 130 amino acid residues and a cytotoxic C-terminal domain of ~ 528 amino acid residues without any known structural homologs [19]. The N-terminal domain of BteA is sufficient to target GFP to lipid rafts of HeLa cell plasma membrane and was therefore termed lipid raft targeting domain (LRT) [19]. Homologous domains were identified by sequence similarity searches in a number of known and putative virulence factors of other bacteria, including a T3SS effector Plu4750 and a MARTX toxin Plu3217 of *Photorhabdus luminescens* which as LRT targeted GFP to lipid rafts [19]. The structure of LRT domain, described by LRT helices (A-A’)-B-C-E, displays an overall tertiary fold similar to 4HBM [20, 21]. Interestingly, the topology of the LRT domain tip is different from that of 4HBM, being formed by the loop region L1, connecting helices A and B, and the capping perpendicular helix D, connecting helices C and E [21]. The *in vivo* membrane-targeting mechanism of LRT domain and its contribution to the cytotoxicity of BteA effector remains poorly characterized. The recombinant LRT domain binds PIP2-containing nanodisks suggesting that BteA may associate with the host plasma membrane and lipid rafts through phospholipid interaction [21]. Interestingly, ectopically expressed BteA of *B. bronchiseptica* remains cytotoxic even upon deletion of the LRT domain or the removal of the first 200 N-terminal amino acids, while its cytotoxicity is diminished upon deletion of the last 14 C-terminal amino acid residues [19, 22].

In this work, we provide a mechanical insight into *in vivo* membrane targeting of the LRT domain and its contribution to BteA cytotoxicity during *B. bronchiseptica* infection.

## Results

### The specificity of the phospholipid binding by LRT motif of BteA effector

To understand the membrane-targeting mechanism of BteA effector, the lipid-binding properties of its N-terminal LRT domain comprising 130 amino acid residues were tested in a protein-lipid overlay assay. The purified recombinant GST-tagged LRT domain of *B. pertussis* BteA (LRT) was used to probe commercial lipid strips. These have been spotted with 100 pmol of different lipids, and bound LRT was detected using an anti-GST antibody. In contrast to GST alone, which displayed no binding affinity for lipids, LRT protein preferentially interacted with negatively-charged lipids, with a preference for phosphatidylinositol 4,5-bisphosphate (PIP2) and phosphatidic acid (PA), as shown in Figs 1A and S1A. The LRT binding depended on the spotted lipid concentration as further corroborated using home-made lipid arrays spotted with a concentration gradient of 10, 100, and 1000 pmol of various lipids or cholesterol per spot (Fig 1B). Interestingly, as also shown in Figs 1A-B and S1A, the GST-tagged full-length BteA protein of *B. pertussis* in complex with its cognate chaperone BtcA (BteA/BtcA) exhibited similar, although, somehow stronger binding of negatively charged lipids than LRT alone. Nevertheless, the lipid-binding abilities of the full-length BteA protein might have been affected by the presence of BtcA, which co-purified with the BteA. In contrast to LRT protein, we were unable to produce soluble BteA without its cognate chaperone BtcA in *Escherichia coli*.

**Fig 1.**
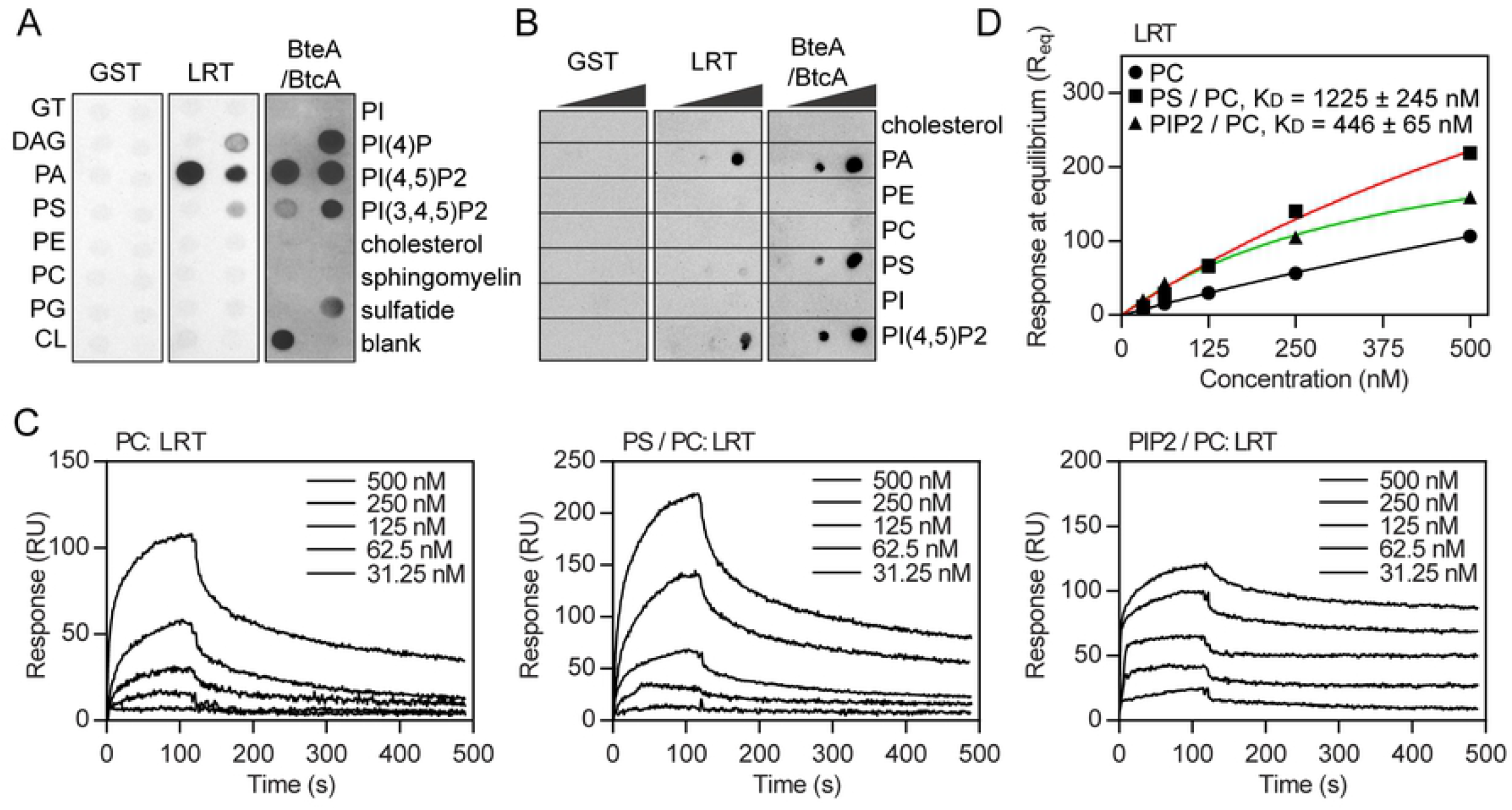
Phospholipid binding by the N-terminal motif of *Bordetella* effector BteA. **(A-B)** Protein-lipid overlay assay. The recombinant GST-tagged N-terminal LRT domain (LRT) and full-length BteA (BteA/BtcA) protein of *B. pertussis* were incubated at 5 μg/ml with commercial **(A)** or home-made **(B)** lipid arrays. The binding was detected using an anti-GST antibody followed by chemiluminescence detection. Recombinant GST was used as a control. Triglyceride (GT); diacylglycerol (DAG); phosphatidic acid (PA); phosphatidylserine (PS); phosphatidylethanolamine (PE); phosphatidylcholine (PC); phosphatidylglycerol (PG); cardiolipin (CL); phosphatidylinositol (PI), and phosphatidylinositol phosphates (PIP, PIP2, PIP3). **(C)** SPR real-time kinetics of LRT binding to lipid membranes. Serially diluted LRT protein (at 500, 250, 125, 62.5 and 31.25 nM concentrations) was injected in parallel over the neutravidin sensor chip coated with the immobilized liposomes (100 nm in diameter) containing PC, PC/PS (80:20) or PC/PIP2 (95:5), and left to associate (120 s) and dissociate (380 s) at constant flow rate of 30 μl/min. The sensograms show the representative binding curves from five independent “one-shot kinetic” experiments. **(D)** SPR steady-state analysis of LRT binding to lipid vesicles. Serially diluted LRT protein was injected over the immobilized lipid vesicles to reach the SPR binding equilibrium, and the near-equilibrium values (R_eq_) were plotted against LRT concentrations (P_0_). Solid lines represent binding isotherms determined by a nonlinear fitting of the data using the equation R_eq_= R_max_/(1 + K_D_/P_0_). The calculated K_D_ values represent the mean ± SD from three independent experiments.

To avoid the limitations of the protein-lipid overlay assay and experiment-to-experiment variability, we next analyzed LRT interaction with a native phospholipid bilayer by surface plasmon resonance (SPR). Three types of large unilamellar vesicles (LUVs) containing phosphatidylcholine (PC) only, or the mixture of PC with phosphatidylserine (PS) and PC with PIP2 at a molar ratio of 80:20 and 95:5, respectively, were prepared and captured on a neutravidin-coated sensor chip. The recombinant proteins were then serially-diluted and injected over the immobilized liposome surface to monitor their binding. While GST alone did not interact with any of the lipid vesicles (S1B Fig), the concentration-dependent binding curves of LRT protein revealed its interaction with all three membrane surfaces with the typical association and dissociation phases of the sensograms (Fig 1C). The binding affinities of LRT to lipid vesicles were next calculated from steady-state binding data as the global fitting of the binding curves to several kinetic models did not provide satisfactory results in terms of χ^2^ and residual statistics. The near-equilibrium values (R_eq_), which were taken from the end of the association phase of the individual binding curves, were plotted as a function of the LRT concentration (Fig 1D), and the equilibrium dissociation constant (K_D_) was determined by nonlinear least-squares analysis of the binding isotherm. As shown in Fig 1D, the binding of LRT to PC vesicles was not saturable (up to 500 nM), indicating a non-specific interaction of the LRT with the charge-neutral phospholipid bilayer. In contrast, the negatively charged lipid surfaces conferred the binding of LRT in a saturable manner, with apparent K_D_ values of 1 225 ± 245 and 446 ± 65 nM for PS- and PIP2-enriched vesicles, respectively. These results, thus, demonstrate that the LRT motif has a direct affinity for negatively charged lipid surfaces *in vitro*.

### Phospholipids PS and PIP2 mediate plasma membrane localization of BteA effector

It was next important to test for the role of PS and PIP2 in guiding the localization of LRT and full-length BteA effector *in vivo*. To this end, the cellular distribution of ectopically expressed GFP-tagged LRT and BteA proteins of *B. pertussis* was analyzed by live-cell fluorescence microscopy in *S. cerevisiae* wild type and mutant strains harboring specific defects in the PS and PIP2 biosynthesis pathways.

As illustrated in Fig 2A, when LRT-GFP was expressed in wild type (WT) strain of *S. cerevisiae* BY4742, it readily associated with the plasma membrane, similarly to its previously reported localization in HeLa cells [19]. The full-length BteA-GFP protein also exhibited peripheral localization in WT strain, although its distribution was patchier (Fig 2A). Upon expression in the *cho1Δ* mutant harboring a deletion of PS synthase Cho1, however, both proteins were distributed throughout the cytoplasm without any plasma membrane localization (Fig 2A). In the same manner, GFP-tagged PS-specific binding protein, GFP-Lact-C2, localized to the plasma membrane of WT strain and dispersed throughout the cytoplasm in the *cho1*Δ mutant (Fig 2A). In contrast, no difference in the localization of PIP2-specific binding protein tagged with GFP, GFP-2xPH(PLCδ), was observed between both strains (S2A Fig). These results demonstrate that both LRT and BteA proteins respond specifically to a decreased level of PS in the *cho1*Δ mutant and point to the importance of PS in their localization.

**Fig 2.**
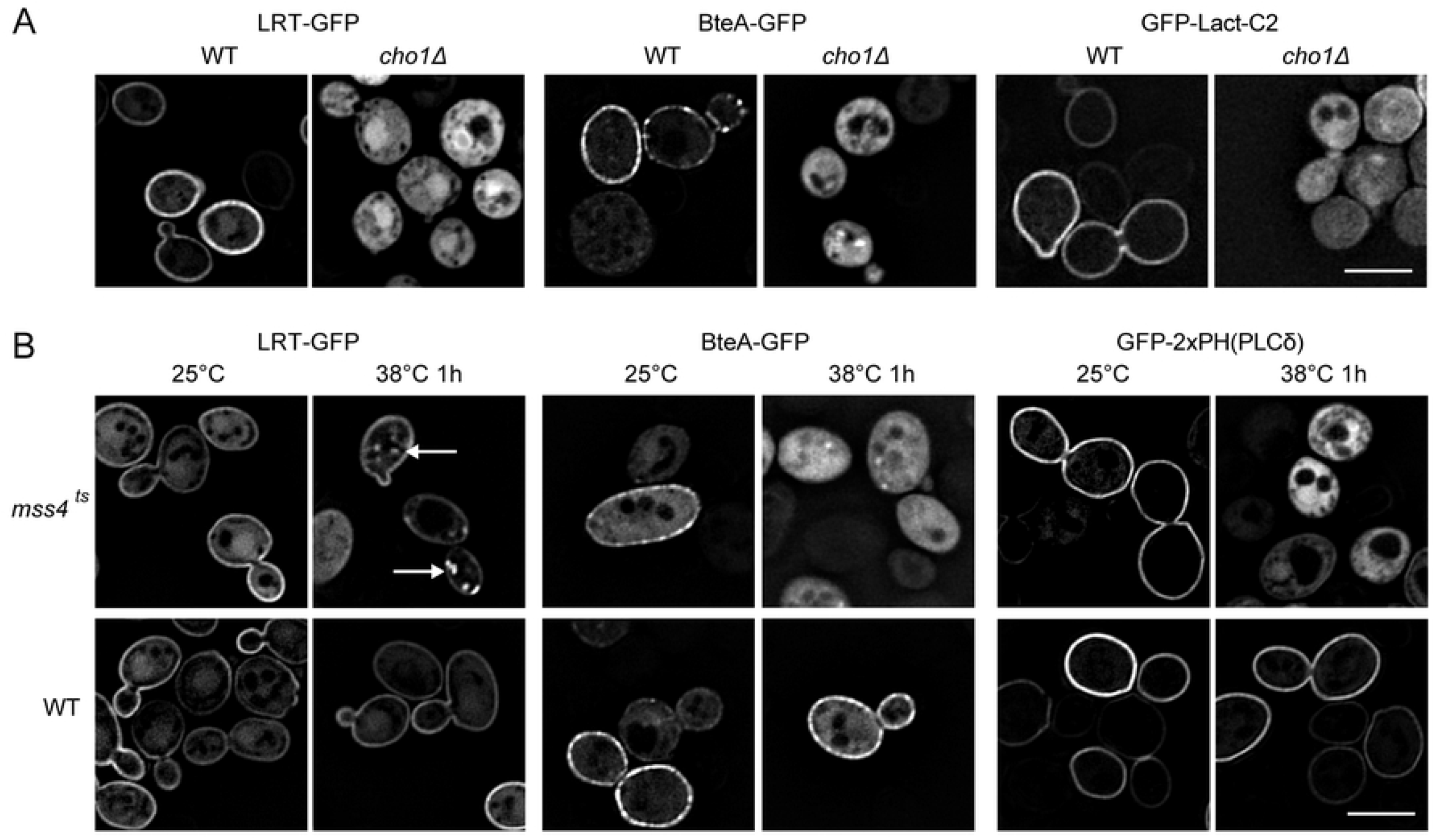
Phospholipids PS and PIP2 guide plasma membrane association of *Bordetella* effector BteA and its LRT motif *in vivo*. Localization of GFP-tagged LRT domain (LRT-GFP) and full-length BteA (BteA-GFP) of *B. pertussis* was analyzed after galactose induction in the following *S. cerevisiae* strains. **(A)** Wild type *S. cerevisiae* BY4742 (WT) and its *cho1Δ* derivative deficient in PS synthesis. **(B)** Wild type *S. cerevisiae* SEY6210 (WT) and its temperature-sensitive *mss4^ts^* derivative exhibiting depleted levels of PIP2 after the shift from permissive (25°C) to restrictive temperature (38°C). White arrows indicate the relocalization of the LRT-GFP protein from the plasma membrane to internal puncta in the *mss4^ts^* mutant. The expression of the GFP-tagged phospholipid probes GFP-Lact-C2 (PS-specific probe) and GFP-2xPH(PLCδ) (PIP2-specific probe) was used to visualize cellular PS and PIP2 levels, respectively. Representative images from two independent experiments with the same outcome are presented. Scale bar, 5 μm.

To test for the role of PIP2, a thermosensitive mutant of the phosphatidylinositol 4-phosphate 5-kinase Mss4 (*mss4*^ts^), which generates PIP2 from PI(4)P was used. To confirm the decreased plasma membrane levels of PIP2 in the *mss4*^ts^ mutant in restrictive temperature, PIP2 levels were visualized by GFP-2xPH(PLCδ). As shown in Fig 2B, GFP-2xPH(PLCδ), indeed, relocated from the plasma membrane to cell cytosol after the shift to restrictive temperature (38°C, 1 hod). In contrast, the localization of PS-specific probe, GFP-Lact-C2, was unaltered, showing that depletion of PIP2 did not merely disrupt the plasma membrane integrity (S2B Fig). As further shown in Fig 2B, LRT-GFP and BteA-GFP proteins localized to the plasma membrane of the *mss4*^ts^ strain at the permissive temperature (25°C), although the association of BteA-GFP was poor when compared to the parental wild type (WT) strain SEY6210 (see S2C Fig for comparison at lower magnification). Importantly, at the restrictive temperature (38°C), BteA-GFP was entirely cytoplasmic-localized (Figs 2B and S2C) while LRT-GFP partially relocated from the plasma membrane to the internal puncta, as highlighted by the white arrows in Fig 2B. Collectively, these data demonstrate that besides PS, also PIP2 levels influence the proper localization of LRT and BteA proteins.

### Definition of structural determinants of the LRT motif in membrane interaction

Having shown that phospholipids guide the localization of BteA effector *in vivo*, we further analyzed the structural determinants of LRT that confer its membrane interaction. The LRT protein bound to lipid vesicles composed of PC only (Fig 1C), suggesting that it associates with lipid membranes at least partly via hydrophobic interactions. Indeed, as depicted in Fig 3A, several hydrophobicity patches, including a protruding hydrophobic leucine 51 (L51) residue, can be visualized on the surface of the LRT structure (aa 29-121, PDB code: 6RGN, [21]). To test for the role of the L51 residue in the binding of LRT to lipid membranes, the residue was replaced by asparagine (L51N) or phenylalanine (L51F), and the capacity of these mutants to interact with lipidic surfaces was evaluated *in vitro* by SPR and *in vivo* by fluorescence microscopy. As shown in Figs 3B and S3A, the substitution of hydrophobic L51 by hydrophilic asparagine decreased the binding of LRT-L51N protein to phospholipid bilayers, regardless of the charge and composition of the immobilized lipid vesicles. In contrast, the replacement of the hydrophobic side chain of L51 by a more hydrophobic aromatic ring of phenylalanine augmented the lipid-binding of LRT-L51F mutant protein. The importance of the L51 hydrophobic side chain in membrane targeting was further corroborated *in vivo* by fluorescence microscopy. As shown in Fig 3C, the L51N variant of LRT-GFP fusion protein was distributed throughout the cytoplasm upon ectopic expression in *S. cerevisiae* cells. In contrast, the wild type protein preferentially localized to the cell periphery. Besides, the introduction of L51F mutation into LRT-GFP fusion protein enhanced its peripheral localization (Fig 3C). Importantly, as shown in S3B Fig by immunoblot analysis, the L51 substitutions within the LRT segment did not affect GFP-fusion protein stability in yeast, and the change of LRT localization was not related to protein degradation. These data, therefore, demonstrate that the hydrophobic moiety of the L51 residue is required for efficient interaction of LRT with the phospholipid bilayer.

**Fig 3.**
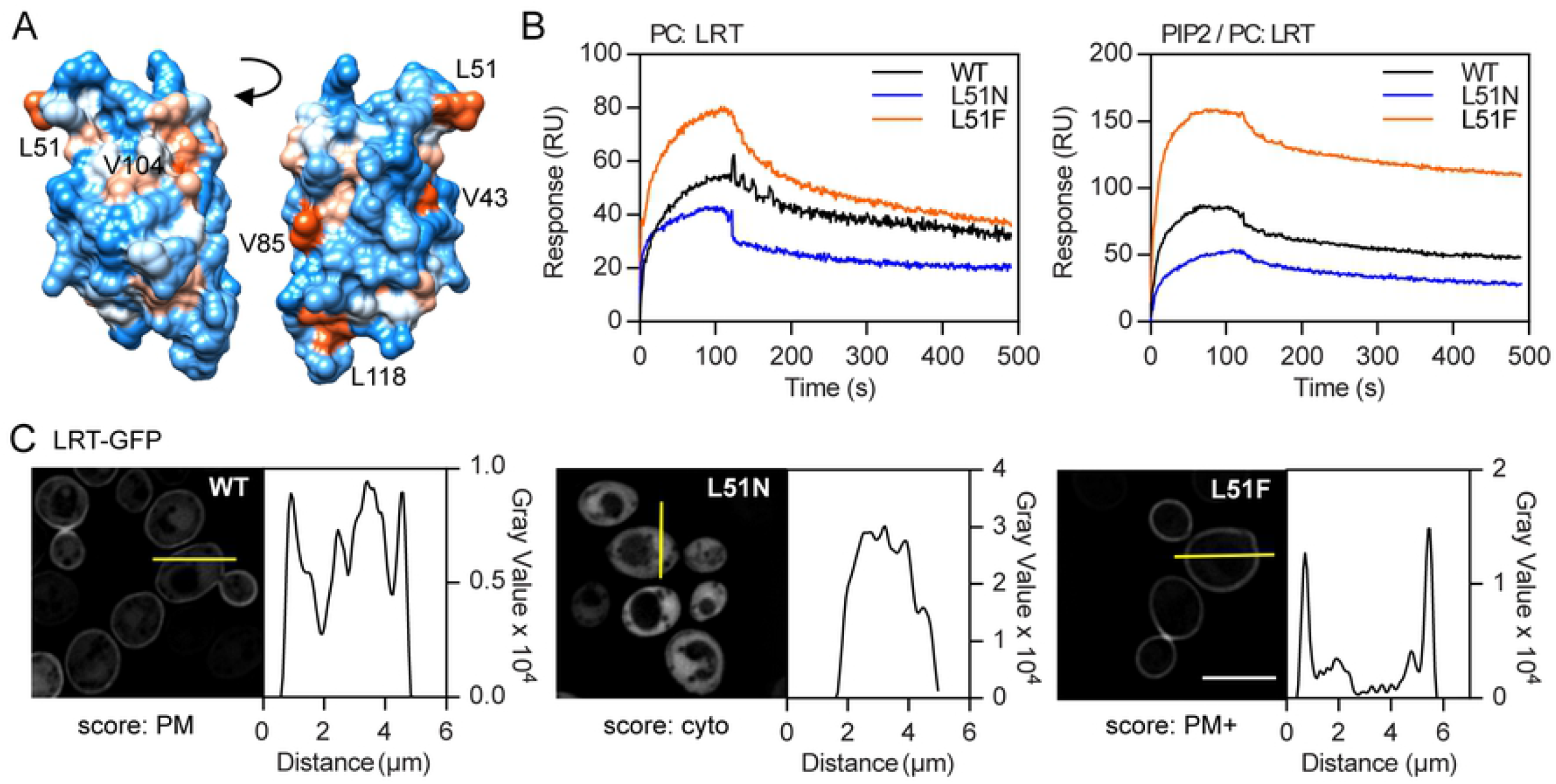
Leu51 residue is involved in hydrophobic interactions of the LRT motif with a phospholipid membrane. **(A)** Surface representation of the LRT structure (aa 29-121, PDB code: 6RGN) colored according to the hydrophobicity and visualized by Chimera 1.14rc. The color gradient ranges from red for the most hydrophobic to white at 0.0 and blue for the most hydrophilic. The exposed hydrophobic residues are indicated. **(B)** Overlay plot of SPR sensograms of the interaction between LRT variants and lipid membranes. Wild type LRT (WT), LRT-L51N, and LRT-L51F proteins at 250 nM concentration were injected over the neutravidin sensor chip coated with the immobilized lipid vesicles containing PC or PC/PIP2 (95:5). The binding curves are representative of five independent “one-shot kinetic” experiments. **(C)** Localization of GFP-fused LRT variants in *S. cerevisiae* BY4741. Yeast cells with plasmids encoding WT or mutant L51N and L51F LRT-GFP fusion proteins were induced for 20 h for protein expression. Representative images from two independent experiments with the same outcome are presented together with a localization score: PM, plasma membrane; Cyto, cytoplasmic, PM+, enhanced PM localization. See Methods for details. Graphs show a fluorescence intensity profile along the yellow bar of a representative cell. Scale bar, 5 μm.

We reasoned that besides LRT-L51 hydrophobic interaction with lipid membrane also electrostatic interactions with negatively charged phospholipid headgroups contribute to membrane binding. Indeed, the structure of LRT revealed a four-helix bundle protein that consists of a large number of positively charged arginine and lysine amino acid residues (Fig 4A-B). To determine critical amino acid residues, a set of LRT mutant proteins harboring charge-reversal substitutions within loop L1 (R50E+H52E+H53E, LRT-L1), helix B (R59E+K62E+R66E, LRT-hB), and helix D (K99E+R100E, LRT-hD) was first constructed, and the capacity of the mutant proteins to interact with the immobilized lipid vesicles was evaluated by SPR. As shown in Fig 4C, the LRT-L1 and LRT-hD mutant proteins exhibited almost complete loss of binding to lipid membranes, regardless of their composition. In contrast, the interaction of LRT-hB with the immobilized PIP2-enriched vesicles was only slightly reduced while completely abolished for the PS-containing liposomes. Hence, the positively charged residues within the loop L1 and helices B and D are required for the interaction with negatively charged membranes, whereas the positively charged side chains at the tip of the LRT structure seem to confer the specificity for PIP2.

**Fig 4.**
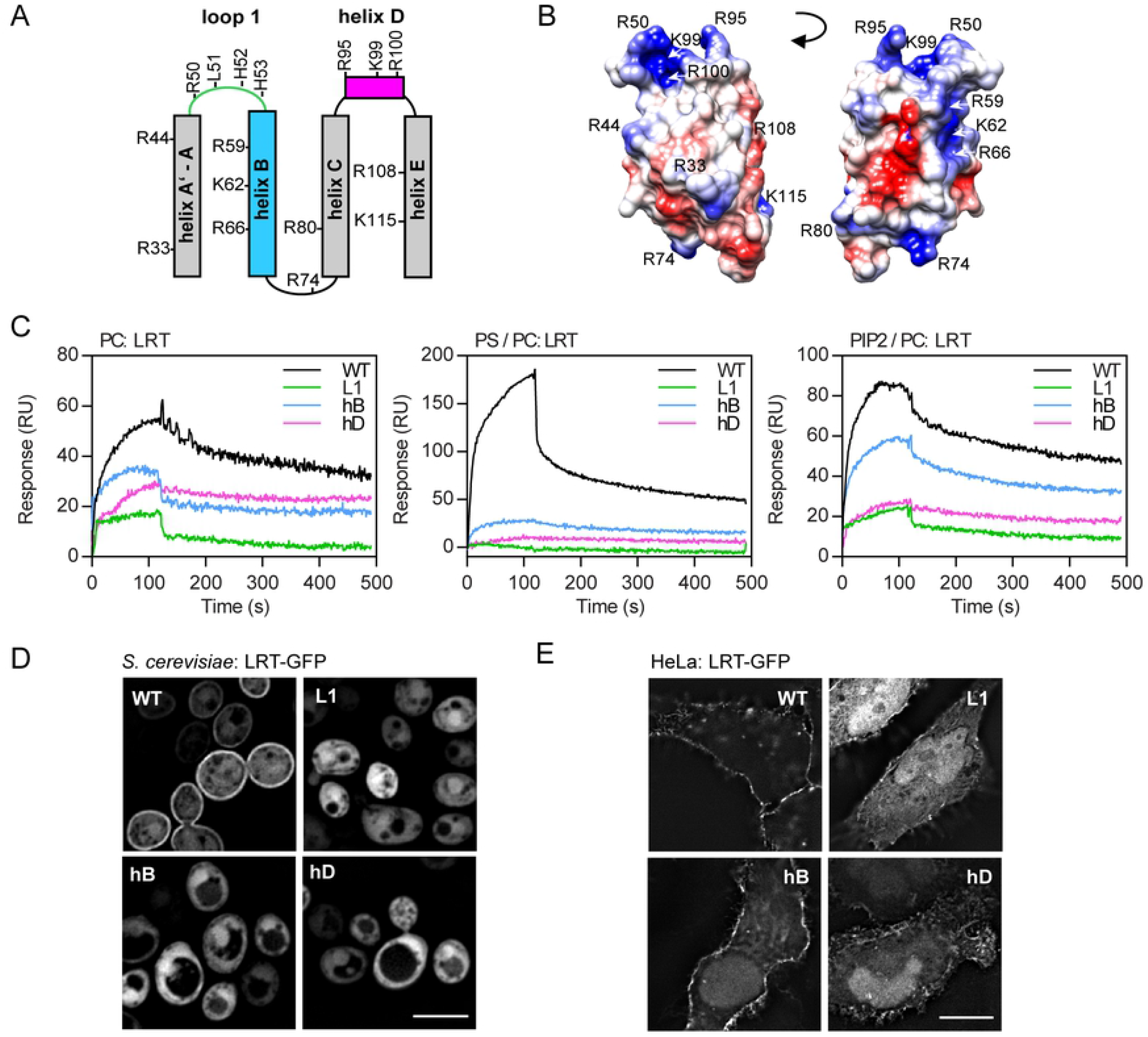
Electrostatic interactions determine the membrane interaction of the LRT domain. **(A-B**) Topology diagram **(A)** and coulombic surface coloring **(B)** of the LRT helix bundle fold (aa 29-121, PDB code: 6RGN) with the indication of positively charged arginine and lysine residues. The color gradient of coulombic surface coloring, as calculated by Chimera 1.14rc, ranges from the positive charge in blue to the negative charge in red. **(C)** Overlay plot of SPR sensograms of the interaction between LRT protein variants and lipid membranes. Wild type LRT (WT) and its charge-reversal substitution L1 (R50E+H52E+H53E), hB (R59E+K62E+R66E) and hD (K99E+R100E) variants were injected at 250 nM concentration over the neutravidin sensor chip coated with the immobilized lipid vesicles containing PC, PC/PS (80:20) or PC/PIP2 (95:5). The sensograms show the representative binding curves obtained from six independent “one-shot kinetic” experiments. **(D-E)** Localization of the GFP-tagged LRT protein variants. **(D)** *S. cerevisiae* BY4741 cells harboring plasmids encoding the indicated LRT-GFP protein variants were induced for 20 h for protein expression and examined by live-cell imaging. Scale bar, 5 μm. **(E)** HeLa cells were transiently transfected with plasmids expressing the indicated GFP-tagged LRT proteins, fixed after 18 h and examined by fluorescence microscopy. Scale bar, 20 μm. Representative images from two independent experiments with the same outcome are presented.

To corroborate this data *in vivo*, localization of mutant GFP-tagged LRT proteins was analyzed by fluorescence microscopy in *S. cerevisiae* and HeLa cells upon LRT ectopic expression. As shown in Fig 4D, charge reversal substitutions within loop L1 (R50E+H52E+H53E, LRT-GFP-L1), helix B (R59E+K62E+R66E, LRT-GFP-hB), and helix D (K99E+R100E, LRT-GFP-hD) resulted in cytoplasmic localization of the mutant proteins in *S. cerevisiae* cells. In transiently transfected HeLa cells, as compared to LRT-GFP, the mutant proteins also predominantly localized to cell cytosol and accumulated in cell nuclei by passive diffusion through nuclear pore complexes (Fig 4E), although slight plasma membrane localization of LRT-GFP-hB was still noticeable. To gain more insight into residue specificity, we further performed glutamic acid and/or alanine mutagenesis screens of arginine, lysine, and histidine residues within the LRT motif. As summarized in Table 1 comprising localization scores for individual mutants and shown by representative micrographs in Figs 5 and S4, substitutions of positively charged amino acid residues within LRT had a rather dramatic impact on membrane association of the GFP-tagged LRT protein in *S. cerevisiae*. Data from both mutagenesis screens revealed that individual residues within loop L1 (H52, H53), helix B (R59, K62), and helix D (R95, K99, and R100) play a critical role in stabilizing membrane interaction of LRT. Besides, residue R108 within helix E was also important.

**Fig 5.**
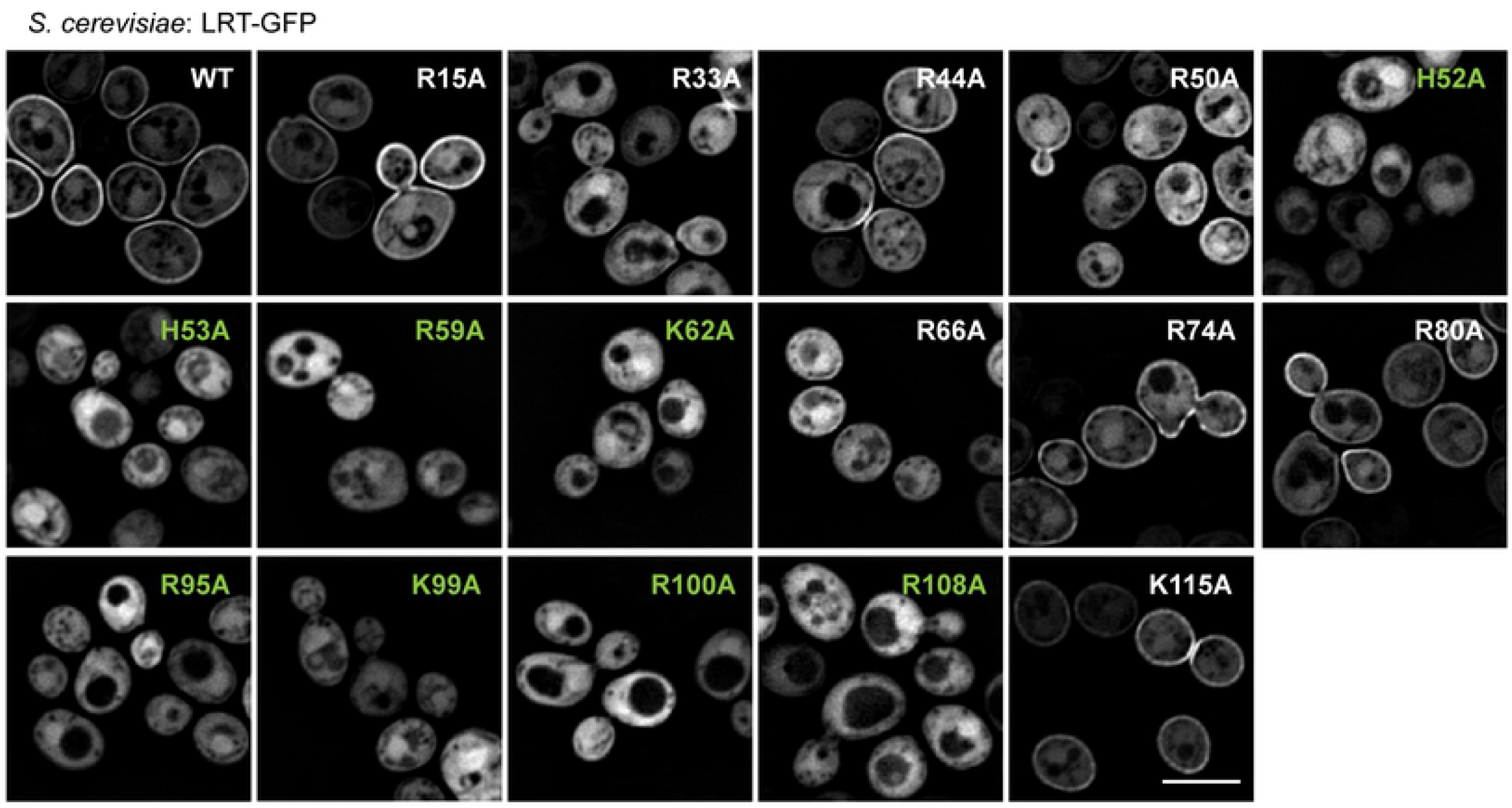
Positively charged residues of the loop L1, helix B, and helix D are critical for the plasma membrane association of the LRT motif. *S. cerevisiae* BY4741 cells carrying plasmids encoding the indicated LRT-GFP protein variants were induced for 20 h for protein expression and examined by live-cell imaging. Representative images from two independent experiments with the same outcome are presented. Scale bar, 5 μm.

**Table 1.** Alanine and glutamic acid mutagenesis of positively charged amino acid residues in the LRT motif of BteA. The cellular distribution of indicated variants of LRT-GFP fusion proteins in *S. cerevisiae* was evaluated using intensity profile plots. The localization score is indicated: PM, plasma membrane; PM-, attenuated PM localization; Cyto, cytoplasmic; ND, not done. See Methods for details. The protein variants with cytoplasmic score are highlighted in green.

| Alanine mutagenesis |  | Glutamic acid mutagenesis |  |
| --- | --- | --- | --- |
| Substitution | Localization score | Substitution | Localization score |
| R15A | PM | R15E | PM |
| R33A | PM- | R33E | Cyto |
| R44A | PM | R44E | PM |
| R50A | PM | R50E | Cyto |
| H52A | Cyto | H52E | ND |
| H53A | Cyto | H53E | ND |
| R59A | Cyto | R59E | Cyto |
| K62A | Cyto | K62E | Cyto |
| R66A | PM- | R66E | PM- |
| R74A | PM | R74E | PM- |
| R80A | PM | R80E | PM |
| R95A | Cyto | R95E | Cyto |
| K99A | Cyto | K99E | Cyto |
| R100A | Cyto | R100E | Cyto |
| R108A | Cyto | R108E | Cyto |
| K115A | PM | K115E | PM- |

### Membrane targeting and phospholipid binding by LRT motif is dispensable for cytotoxic activity of BteA

To get a better insight into the targeting and cytotoxicity mechanisms of BteA, we next introduced the charge-reversal substitution of L1 (R50E+H52E+H53E) into the full-length protein and tested the abilities of the resulting BteA-L1 mutant to bind membrane phospholipids, localize to the plasma membrane and induce cellular cytotoxicity.

The purified mutant BteA-L1 protein in complex with BtcA (BteA-L1/BtcA) was unable to bind any of the negatively charged lipids in contrast to the wild type BteA/BtcA complex, binding PA, PS, and PIP2, as determined by protein-lipid overlay assay and shown in Fig 6A. Furthermore, the ectopically expressed mutant GFP-tagged BteA (BteA-L1) of *B. pertussis* was distributed throughout the cytoplasm of *S. cerevisiae*, whereas the wild type protein localized to the cell plasma membrane (Fig 6B). We were unable to visualize the full-length BteA protein of *B. bronchiseptica* in *S. cerevisiae* or BteA proteins of *B. pertussis* and *B. bronchiseptica* in transiently-transfected HeLa cells due to the very high toxicity of these proteins (data not shown), as previously reported [18, 19]. However, both GFP-tagged mutants of *B. pertussis* and *B. bronchiseptica* BteA without the last 14 residues (BteA^1-642^-L1 and *Bb*BteA^1-644^-L1, respectively) localized to cell cytosol in transiently-transfected HeLa cells (Fig 6C). In contrast, the wild type variants of these truncated proteins targeted cell plasma membrane, as also shown in Fig 6C. Taken together, our data showed that charge-reversal substitution within loop L1 of LRT disrupted the lipid binding and membrane localization also for the full-length BteA protein.

**Fig 6.**
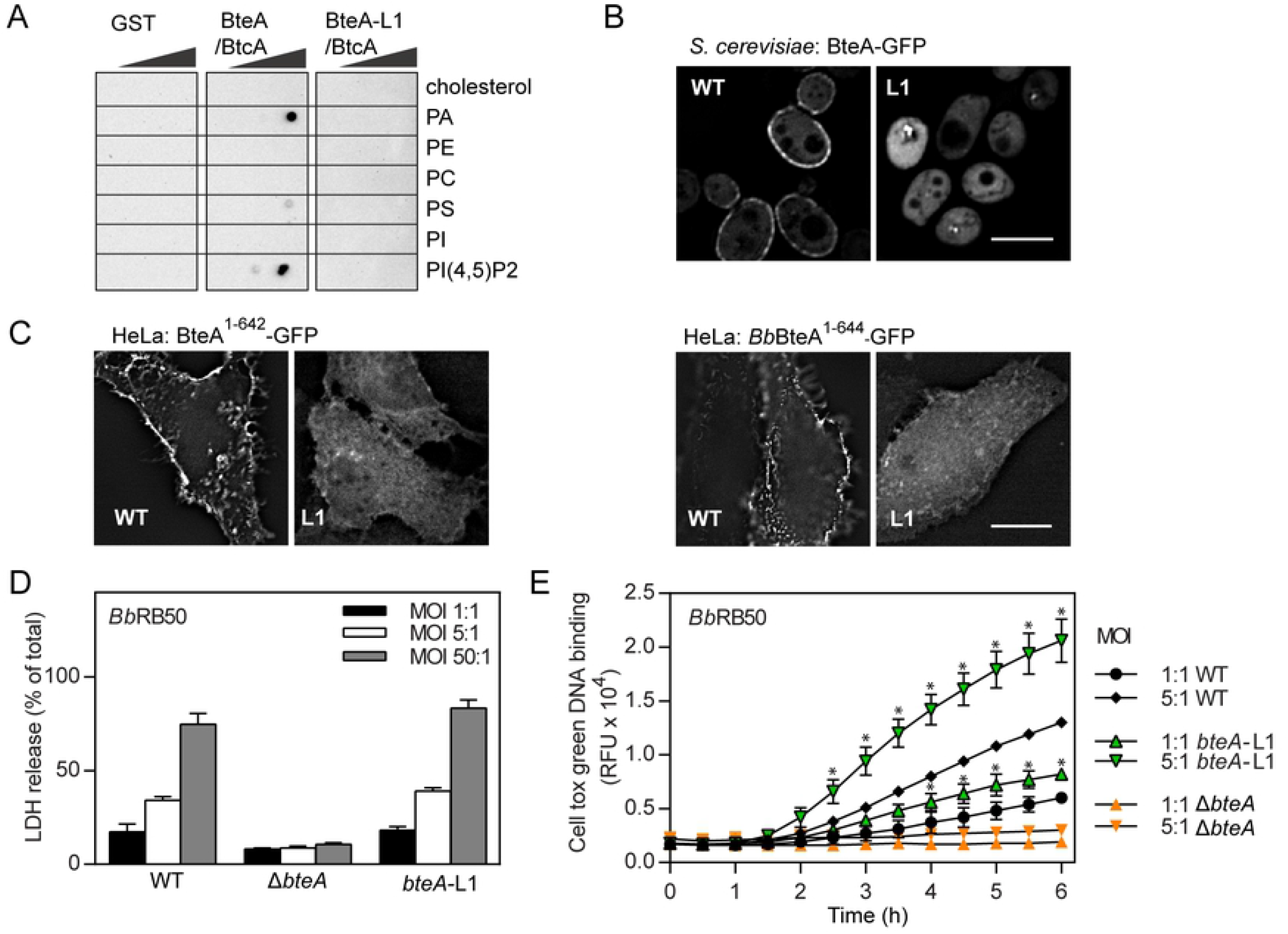
Plasma membrane targeting of BteA effector does not contribute to its cytotoxicity. **(A)** Protein-lipid overlay assay. Recombinant GST-tagged full-length *B. pertussis* BteA (BteA/BtcA) and its mutated variant with charge reversal substitutions within the loop L1 (BteA-L1/BtcA; R50E+H52E+H53E) were incubated with home-made lipid arrays at 5 μg/ml. The BteA binding was detected using an anti-GST antibody, followed by chemiluminescence detection. Recombinant GST was used as a control. See the legend of Fig 1B for the description of spotted lipids. Results are representative of two independent experiments. **(B-C)** Localization of the GFP-tagged BteA protein variants. **(B)** *S. cerevisiae* BY4741 cells harboring plasmids encoding GFP-tagged full-length BteA of *B. pertussis* and its variant BteA-GFP-L1 (L1; R50E+H52E+H53E) were cultivated for 20 h in the medium supplemented with galactose to induce protein expression. Scale bar, 5 μm. **(C)** HeLa cells were transiently transfected with plasmids expressing GFP-tagged variants of *B. pertussis* BteA (BteA-1-642-GFP, BteA-1-642-L1-GFP) or *B. bronchiseptica* BteA (*Bb*BteA-1-644-GFP, *Bb*BteA-1-644-L1-GFP). After 18 h, the HeLa cells were fixed and examined by fluorescence microscopy. Scale bar, 20 μm. Representative images from two independent experiments with the same outcome are presented. **(D-E)** HeLa cells were infected with WT, *ΔbteA*, and *bteA*-L1 mutant derivatives of *B. bronchiseptica* RB50 at the indicated multiplicity of infection (MOI). Cytotoxicity was measured as lactate dehydrogenase (LDH) release 3 h post-infection (**D**) and as real-time kinetics of membrane permeabilization determined by fluorescent DNA binding dye CellTox Green **(E)**. Values represent the means ± SE from 2 independent experiments performed in duplicate (n = 4). * p < 0.05, bteA-L1 vs. WT-infected cells, unpaired two-tailed t-test using the corresponding infection time and MOI.

To determine the role of plasma membrane targeting in BteA-induced cytotoxicity, the charge-reversal substitution within loop L1 (R50E+H52E+H53E, L1) was next introduced into the *bteA* open reading frame on the chromosome of *B. bronchiseptica* RB50, and the cytotoxic capacity of the mutant bacteria was tested. As shown in Fig 6D, the cytotoxic potency of the *bteA*-L1 mutant bacteria measured as release of the intracellular enzyme lactate dehydrogenase (LDH) into cell culture media was comparable to that of the wild type strain. After 3 h of infection, both strains induced the lysis of 40 % and 80 % of HeLa cells at MOI 5:1 and 50:1, respectively. In contrast, no cell lysis was provoked by infection with Δ*bteA* strain, harboring an in-frame deletion of the BteA effector. To assess HeLa cell cytotoxicity during *B. bronchiseptica* infection in a more sensitive way, we also monitored the real-time kinetics of membrane permeabilization using a fluorescent DNA binding dye CellTox Green. Intriguingly, infection of HeLa cells by the *bteA-L1* mutant bacteria at MOI 1:1 and 5:1 appeared to disrupt membrane integrity substantially more than the wild type bacteria at the same MOI (Fig 6E). Again, no membrane permeabilization was provoked by *ΔbteA* mutant strain, showing that all observed cytotoxicity of the RB50 towards HeLa cells was due to the action of the BteA effector. Taken together, these data revealed that the LRT domain does not promote BteA cytotoxicity by targeting BteA to plasma membrane of the host cells.

## Discussion

In an attempt to decipher the mechanism by which the LRT domain of BteA effector targets the plasma membrane, we report here that recombinant LRT protein binds negatively-charged membrane phospholipids with a preference for PIP2. Besides, we show that the domain ability to bind phospholipids and interact with membranes *in vitro* is necessary for its plasma membrane targeting in yeast and HeLa cells. Moreover, the depletion of PS and PIP2 in yeast cells mislocalizes the GFP-tagged LRT domain and full-length BteA from the plasma membrane. This data shows that the association of BteA effector with the plasma membrane is guided by phospholipids PS and PIP2 *in vivo*. We also demonstrate that LRT-mediated lipid binding and plasma membrane targeting are dispensable for the cytotoxic activity of BteA effector.

Using a protein-lipid overlay assay, we showed that recombinant LRT protein encoding the membrane localization domain of BteA binds to the negatively-charged phospholipids. The domain binds preferentially to PIP2 (a.k.a PI(4,5)P2) and PI(3,4)P2, as compared to PI(3,5)P2 regioisomer or various other phosphatidylinositol phosphates PI(3)P, P(4)P, P(5)P (*c.f*. S1A Fig). This data goes well with a recent biophysical study, which detected LRT binding to PIP2-containing lipoprotein nanodiscs by shifts of NMR cross-peaks and performed NMR titration experiments using soluble phosphatidylinositol analogs [21]. The lipid-binding site of the LRT domain may thus favor the presence of two phosphate groups on the inositol ring next to each other. LRT protein, however, also binds to negatively-charged lipids, such as PA or PS. Nevertheless, the LRT affinity for PS-containing liposomes was about 2.7 fold lower (Kd ~1.2 μM) than for those enriched in PIP2, exhibiting Kd of ~450 nM. The affinities of the LRT domain for lipids are well comparable to other lipid-binding domains. For example, the eukaryotic PH domain of PLCδ binds to PIP2 with Kd of ~ 2 μM [23] while binding of full-length *Pseudomonas aeruginosa* ExoU effector to PIP2-enriched liposomes is characterized by Kd of ~110 nM [24]. Unfortunately, we could not estimate binding affinities of full-length recombinant BteA to PIP2- and PS-enriched liposomes as soluble BteA did not purify without its chaperone BtcA. This cognate chaperone binds to the LRT domain and escorts the effector for the type III secretion in *Bordetella* [25].

The surface of the LRT domain structure displays a positively charged interface, which is formed by loop L1, helix B, and helix D (*c.f*. Fig 4B). A similar positively charged interface formed by L1 and L3 at the domain tip is present within the 4HBM domain family identified in diverse bacterial toxins [7]. The 4HBM domains associate with membranes through positively charged residues of this interface in cooperation with surface-exposed hydrophobic residue at the top of loop L1 [7, 8]. Importantly, surface analysis of the LRT structure pointed out an exposed hydrophobic L51 residue within loop L1 that aligns with hydrophobic residue at position 16 or 17 of 4HBM, as shown in Fig 7A. The L51 residue is surrounded by several basic residues, including H52+H53, which align with conserved K/R residue at position 18 of 4HBM (Fig 7A). Moreover, the substitution of the L51 residue with hydrophilic asparagine (L51N) resulted in reduced membrane binding and plasma membrane association, whereas substitution with more hydrophobic phenylalanine (L51F) increased membrane binding and plasma membrane targeting of the mutant domain variant (*c.f*. Fig 3). Besides, extensive mutagenesis of LRT residues revealed that substitutions of basic residues at the domain tip or in its proximity disrupted plasma membrane targeting of the mutant LRT variants in *S. cerevisiae*. In contrast, substitutions distant from the domain tip showed no alteration (*c.f*. Table 1). Our experimental data and model obtained by molecular docking of PIP2 headgroups to LRT structure (Fig 7B), hence, suggest that membrane targeting by LRT is reminiscent of that of 4HBM. The L51 residue of the L1 region may directly penetrate the apolar membrane milieu, whereas the positively charged residues at the domain tip and helix B would provide a combination of electrostatic attraction to the negatively charged membrane surface and a specific headgroup recognition of the membrane phospholipids (Fig 7B). Besides, helix D of LRT (Fig 7B), which is analogous to L3 of 4HBM, may provide structure-stabilizing interactions. Indeed, the critical residue R100 within helix D aligns with conserved R71 residue of 4HBM and is positioned towards the structure interior. We, therefore, propose that the LRT domain of *Bordetella* BteA constitutes an additional member of 4HBM family of bacterial proteins that targets host plasma membrane by a basic-hydrophobic motif [7, 8]. The proposed model of LRT interaction characterized by the membrane-penetrating leucine residue within loop L1 differs from the “side-on” interaction model suggested within the recent biophysical study [21]. However, similarly to this study, we identified the critical role of residues R59 and K62 within helix B for membrane interaction of LRT (Fig. 7B). Further work will be needed to address the membrane targeting of homologous LRT-like domains. Importantly, we were able to locate a homologous hydrophobic leucine or isoleucine residue within the L1 loops of these domains.

**Fig 7.**
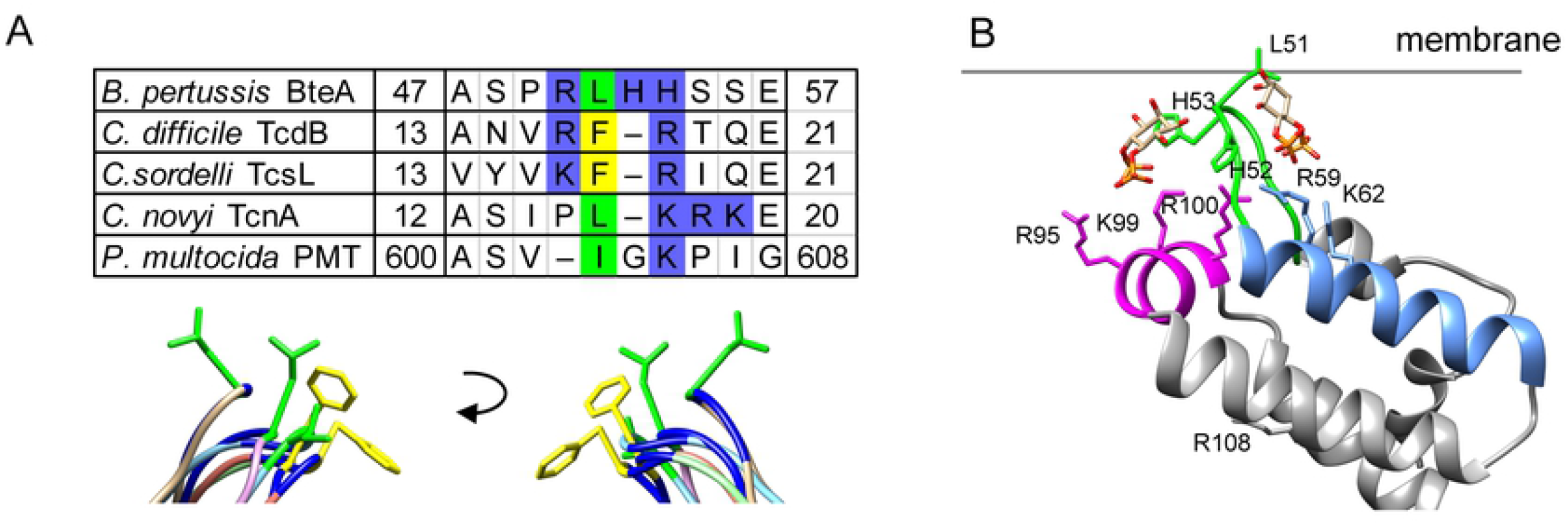
Comparison of LRT with related PDB structures and its membrane association model. **(A)** Amino acid and structure alignments of the L1 region. Amino acids I/L are highlighted in green, F are highlighted in yellow, H/K/R are highlighted in blue. The structural alignment of the LRT domain of *B. pertussis* BteA (brown, PDB code: 6RGN) with 4HBM domains of *C. difficile* TcdB (orange, PDB code: 5UQM), *C. sordelii* TcsL (light green, PDB code: 2VKD), *C. novyi* TcnA (pink, PDB code: 2VK9), and *P. multocida* PMT (light blue, PDB code: 2EBF) was performed using Chimera 1.14rc. **(B)** Model of membrane association of the LRT domain. Critical amino acid residues identified by domain mutagenesis are depicted as sticks within the ribbon representation of the LRT structure and were visualized by Chimera 1.14rc. The headgroup of PIP2 (inositol 4,5-bisphosphate) was docked by AutoDock Vina. The binding energy was almost equal for both ligand binding sites, exhibiting the binding affinity value of −3.7 kcal/mol.

The intriguing question is the differential specificity of the LRT domain of BteA as compared to the 4HBM family and the reasons for its previously reported capacity to target lipid rafts in mammalian cells [19]. *In vivo*, both PIP2 and PS phospholipids were critical for plasma membrane targeting of GFP-tagged LRT domain and BteA effector in yeast cells, the model that can survive intracellular expression of the full-length BteA protein of *B. pertussis* [18]. Indeed, PIP2, which constitutes only about 0.05% of the eukaryotic cell phospholipids, is highly enriched in the cytosolic leaflet of the plasma membrane (~1 % of plasma membrane phospholipids) where it controls many cell processes, including vesicle trafficking, ion channel modulation, and actin-cytoskeleton dynamics [26-28]. Furthermore, depending on the cell type and experimental conditions, enrichment of PIP2 in microdomains of various sizes and/or detergent-insoluble domains has been reported [29-32]. PS, on the other hand, is the main negatively charged phospholipid present in the plasma membrane of eukaryotic cells, comprising ~ 34% of plasma membrane phospholipids in *S. cerevisiae* and ~ 8% of those in mammalian cells [33]. Interestingly, PIP2 and PS were reported to have polarized localization in the yeast cells, accumulating in bud necks and the bud cortex [34]. It is, therefore, worth mentioning that we have noticed polarized localization of GFP-tagged LRT domain and full-length BteA during live-cell imaging of yeast cells, as shown in S5 Fig. The LRT domain exhibited continuous plasma membrane distribution and accumulated at the incipient bud site, small bud, and at the latter stages of the cell cycle to the mother-bud neck. The distribution of full-length BteA did not oscillate so highly during the cell cycle and was more patch-like, suggesting additional interactions of BteA at the cell periphery as compared to the LRT domain (S5 Fig). Furthermore, in contrast to LRT, BteA displayed difficulties to localize to plasma membrane in the *mss4^ts^* mutant even at permissive conditions (Fig 2B). In these conditions, *mss4*^ts^ cells have 2-fold lower levels of PIP2 as compared to the wild type cells [35] and exhibit disturbance of their secretory pathways and/or actin cytoskeleton [36]. This might have affected the delivery and localization of the BteA effector and/or its putative interactor. Nevertheless, despite other putative interactions of BteA, its plasma membrane targeting was entirely dependent on the LRT domain and abolished upon charge-reversal substitution within loop L1 of LRT *(c.f*. Fig 6). It remains to be established whether the LRT domain has one or more additional interacting partners other than PIP2 and PS that would further shape its specificity for lipid rafts in mammalian cells.

The mechanism of cytotoxic activity of BteA inside host cells remains unknown. Previous experiments have shown that BteA construct lacking the LRT domain displayed cell cytotoxicity indistinguishable from the full-length BteA following transient transfection [19]. However, the need for LRT-dependent localization in these experiments might have been masked by the expression of BteA at high levels. By introduction of charge-reversal substitution within loop L1 of LRT into the *bteA* open reading frame on the chromosome of *B. bronchiseptica* RB50 we found here that LRT-mediated targeting to plasma membrane does not facilitate the cytotoxic activity of BteA during bacterial infection (*c.f*. Fig 6D-E). Moreover, at low MOI, bacteria encoding mutant BteA-L1 caused significantly more membrane permeabilization than the wild type bacteria (*c.f*. Fig 6E). This data contrasts with the requirement of membrane targeting in cytotoxicity of 4HBM-comprising bacterial toxins that have a membrane-localized target. For example, the impaired membrane localization of the *C. sordellii* TcsL mutants F17N and R18A (L1 mutations) correlated with a decrease in TscL cytotoxicity [10]. Our data, hence, suggest that BteA induces cytotoxicity by a cellular pathway also initiated from the host cytoplasm. It remains to be established what is the purpose of plasma membrane localization of BteA and what kind of targets of BteA are present there.

## Methods

### Bacterial and yeast strains, cell lines, media, and plasmids

All bacterial and yeast strains used in this study are listed in S1 and S2 Tables, respectively, whereas the plasmids are listed in S3 Table. The *E. coli* strain XL1-Blue was used for plasmid construction, *E. coli* strain Rosetta 2 was employed for recombinant protein production, and *E. coli* strain SM10 λpir was used for plasmid transfer into *Bordetella* by bacterial conjugation. *E. coli* strains were grown on Luria-Bertani (LB) agar medium or in LB broth, and when appropriate, media were supplemented with 100 μg/ml ampicillin and/or 30 μg/ml kanamycin. *Bordetella bronchiseptica* strains were grown on Bordet-Gengou (BG) agar medium (Difco, USA) supplemented with 1% glycerol and 15% defibrinated sheep blood (LabMediaServis, Jaromer, Czech Republic) at 37°C and 5% CO2, or in modified Stainer-Scholte (SSM) medium supplemented with 5 g/l of Casamino Acids at 37°C. Mutant *B. bronchiseptica* strains were constructed by homologous recombination using the suicide allelic exchange vector pSS4245, as described previously [18]. The modified portions of the *B. bronchiseptica* chromosome were verified by sequencing (Eurofins Genomics, Germany). *Saccharomyces cerevisiae* strains were grown in standard rich medium (YPD) or synthetic defined medium supplemented with appropriate amino acids and 2% glucose (SD media, non-inducing media) or 2% galactose (SC-galactose media, inducing media). The media for *S. cerevisiae* BY4742 *cho1Δ* were further supplemented with 1 mM ethanolamine (Sigma Aldrich, USA). The *S. cerevisiae* SEY6210 strains, wild type (WT), and the *mss4^ts^-102 (mss4^ts^)* mutant were generous gifts of Chris Stefan (London, UK). Cell lines HeLa (ATCC CCL-2, human cervical adenocarcinoma) was cultivated in Dulbecco’s Modified Eagle Medium (DMEM) supplemented with 10% (vol/vol) heat-inactivated fetal bovine serum (FBS) at 37 °C and 5% CO2. The plasmids used in the study were constructed using T4 DNA ligase, or the Gibson assembly strategy. PCR amplifications were performed using Herculase II Phusion DNA polymerase (Agilent, USA) from chromosomal DNA of *B. pertussis* B1917 or *B. bronchiseptica* RB50, and PCR mutagenesis was employed to introduce the site-directed substitutions within the membrane localization domain of BteA. All constructs were verified by DNA sequencing (Eurofins Genomics, Germany). The plasmid pRS426GFP-2xPH(PLCδ) encoding the yeast PIP2-specific probe GFP-2xPH(PLCδ) [35] was a kind gift from Chris Stefan (London, UK), and the PS-specific probe GFP-Lact-C2 encoded by pGPD416-GFP-Lact-C2 [34] was generously provided by Vanina Zaremberg (Calgary, Canada).

### Production and purification of recombinant BteA proteins

The recombinant proteins control GST, and GST-tagged wild type and mutated variants of the full-length BteA protein (BteA, aa 1-656) of *B. pertussis* B1917 and its N-terminal membrane localization domain (LRT, aa 1-130) were produced in *E. coli* Rosetta 2 from pGEX-6P1 expression vector (GE Healthcare, USA). To improve the solubility of the full-length protein, BteA was produced concomitantly with its chaperone BtcA expressed from the pET28b vector (Novagen, USA) (S3 Table). Exponential *E. coli* cultures grown at 30°C were induced for protein production by the addition of IPTG to 0.1 mM at OD_600_ = 0.3 and grown for an additional 16 h at 20 °C. Bacterial cells were harvested by centrifugation, and the cell pellet was resuspended in ice-cold 50 mM Tris-HCl pH 7.4, 150 mM NaCl, and Complete Mini protease inhibitors (EDTA free, Roche, Switzerland). Bacterial cells were disrupted by ultrasound, and the lysate was clarified by centrifugation (20,000 g, 30 min). The recombinant proteins were purified from the supernatant fraction using columns prepacked with Glutathione-Sepharose 4B (Amersham, UK). The resin with bound proteins was washed with 50 mM Tris-HCl pH 7.4, 150 mM NaCl and proteins were eluted with 10 mM reduced glutathione in 50 mM Tris-HCl pH 7.4, 150 mM NaCl. Protein preparations were dialyzed overnight into 50 mM Tris-HCl pH 7.4 and 150 mM NaCl. The integrity and purity of recombinant proteins were verified by SDS-PAGE electrophoresis followed by Coomassie blue staining, and protein concentration was determined by the Bradford protein assay.

### Protein-lipid overlay assay

Home-made lipid arrays and commercial lipid strips (cat.no. P-6002, membrane lipid strip, and cat.no. P-6001, PIP strip, both Echelon Biosciences, USA) were used to test lipid binding of recombinant LRT and BteA proteins. To prepare home-made lipid arrays, solutions of 5 μM, 50 μM and 500 μM of cholesterol (cat.no.C8667, Sigma Aldrich, USA), phosphatidic acid (PA 16:0-18:1; cat.no.840857P, Avanti Polar Lipids, USA,), phosphatidylethanolamine (PE 16:0-18:1; cat.no.01991, Sigma Aldrich, USA), phosphatidylcholine (PC 16:0-18:1; cat.no.42773, Sigma Aldrich, USA), phosphatidylserine (PS 16:0-18:1; cat.no.840034P, Avanti Polar Lipids, USA), phosphatidylinositol (PI 16:0-18:1; cat.no.850142P, Avanti Polar Lipids, USA) or phosphatidylinositol 4,5-bisphosphate (PI(4,5)P2 18:1-18:1; cat.no.850155P, Avanti Polar Lipids, USA) were prepared in chloroform or in a 20:9:1 mixture of chloroform / methanol / H_2_O in the case of PIP2. Using a Hamilton syringe, 2 μl of each concentration was then spotted on the nitrocellulose membrane (BioTrace NT, Pall Corporation, USA), yielding 10, 100, or 1000 pmol of cholesterol or lipid per spot. To analyze recombinant protein binding, the home-made arrays or commercial lipid strips (100 pmol of lipid per spot) were blocked for 1 h with PBS/0.1% (v/v) Tween-20 (PBST) containing 3% (w/v) BSA (PBST-BSA) at room temperature (RT) prior to 1 h-incubation with control GST (5 μg/ml) or recombinant GST-BteA protein derivatives (5 μg/ml) in PBST-BSA. Membranes were then washed with PBST (3x 5 min), probed for 1 h with rabbit anti-GST antibody (1:2,000; clone 91G1, Cell Signaling Technology, USA) in PBST-BSA, and washed again with PBST (3x 5 min). The membranes were further incubated with horseradish peroxidase (HRP)-conjugated anti-rabbit IgG secondary antibody (1:3,000; GE Healthcare, USA) in PBST-BSA for 1 h. The chemiluminescence signal was detected using a Pierce ECL chemiluminescence substrate (Thermo Fisher Scientific, USA) and an Image Quant LAS 4000 station (GE Healthcare, USA).

### Liposome preparation

Large unilamellar liposomes/vesicles (LUVs) were prepared by the thin-film hydration method as previously described by [37]. In brief, phosphatidylcholine (PC 16:0-18:1; cat.no. 42773, Sigma Aldrich, USA), or a mixture of PC with phosphatidylserine (PS 16:0-18:1; cat.no. 840034P, Avanti Polar Lipids, USA) at a molar ratio of 80:20 (PC:PS), and PC with phosphatidylinositol 4,5-bisphosphate (PI(4,5)P2 18:1-18:1; cat.no. 850155P, Polar Lipids, USA,) at a molar ratio of 95:5 (PC:PIP2) were dissolved in chloroform and dried in 10×50 mm glass test tube by rotary evaporation under a nitrogen stream. The residual solvent was removed under high vacuum for 1 h. The lipid film was hydrated at lipid:DNA ratio 1200:1 with 50 mM Tris-HCl pH 7.4, 150 mM NaCl buffer supplemented with an oligonucleotide of the sequence 5’-TATTTCTGATGTCCACCCCC-3’ modified at the 3’ end with cholesterol (Generi-Biotech, Czech Republic). The unilamellar vesicles were prepared by extrusion of the hydrated lipid mixture through a 0.1 μm Whatman Nuclepore Track-Etched polycarbonate membrane (Sigma, USA) assembled in a LiposoFast Basic apparatus (Avestin, Canada). The liposome suspension was supplemented with 8 mM anti-sense oligonucleotide of the sequence 5’-TGGACATCAGAAATACCCCC-3’ modified at the 3’ end by biotin (Generi-Biotech, Czech Republic), and the mixture was centrifuged at 40,000 g for 30 min at 4 °C to remove free biotinylated ssDNA. The pelleted liposomes were resuspended in 50 mM Tris-HCl pH 7.4, 150 mM NaCl, and used for binding experiments.

### Surface plasmon resonance (SPR)

The SPR experiments were performed on ProteOn XPR36 Protein Interaction Array System (Bio-Rad, USA) at 25 °C as previously described by [37]. In brief, lipid vesicles (100 μg/ml) were immobilized to a neutravidin-coated NLC chip (Bio-Rad, USA) at a flow rate of 30 μl/min to a coupling level of about 1000 resonance units (RU). The proteins, as indicated in the figure legends, were serially diluted in the SPR running buffer, containing 50 mM Tris-HCl pH 7.4, 150 mM NaCl and 0.005% Tween-20, and injected in parallel over the lipid surface at a constant flow rate of 30 μl/min (‘one-shot kinetics’). The sensograms were corrected for sensor background by interspot referencing (the sites within the 6 × 6 array that were not exposed to ligand immobilization but were exposed to analyte flow) and double referenced by subtraction of the analyte using a “blank” injection. Assuming a Langmuir-type binding between the protein (P) and protein binding sites (S) on vesicles (i.e., P + S ↔ PS), near-equilibration (R_eq_) values were then plotted against protein concentration (P_0_), and the K_D_ value was determined by nonlinear least-squares analysis of the binding isotherm using the equation R_eq_ = R_max_ / (1 + K_D_/P_0_).

### Yeast cell handling and live-cell fluorescence microscopy

Yeast cell transformation was carried out according to the one-step protocol of Chen *et al*. [38]. The yeast centromeric vector pYC2-CT (Invitrogen, USA) was used for galactose-inducible expression of the BteA protein variants harboring the GFP tag on their C-terminus (S3 Table). The production of BteA-derivative proteins was induced for 20 h by cultivation in SC-galactose inducing media. For depletion of PIP2 from the plasma membrane in *mss4^ts^* strain, cells were incubated at the restrictive temperature (38 °C) for 1 h. To reduce the background fluorescence during live-cell imaging, yeast cells were washed in low-fluorescence synthetic-complete medium (LF-SC) and mounted by covering with a thin slice of 1.5% agarose prepared in LF-SC on a microscope cover glass. Widefield microscopy was performed using an Olympus IX-81 inverted microscope with 100x PlanApochromat oil-immersion objective (N.A. = 1.4) and a Hamamatsu Orca-ER-1394 digital camera. GFP fluorescence was detected with the filter block U-MGFPHQ, exc. 460-488 nm, em. 495-540 nm. Z-stacks were taken with 0.28 μm z-steps, and images were collected in a 16-bit format. Deconvolution with Advanced Maximum Likelihood (AMLE) filters (Xcellence Imaging SW, Olympus) and processing using Olympus Cell-R Xcellence and FIJI (ImageJ) [39] software packages were performed. Cropped images were adjusted for brightness/contrast only and mounted in Adobe Illustrator (Adobe, USA). A single focal plane of a Z-stack is presented in all figures.

### Determination of localization score

The ability of each LRT-GFP variant to associate with yeast plasma membrane was determined by scoring the fluorescence intensity distribution across individual cells in FIJI (ImageJ) [39]. The cells displaying similar levels of fluorescence were selected in the acquired images of different LRT-GFP variants, and the “Line tool” of FIJI was used to draw the line across individual cells. The line was drawn such to avoid yeast vacuole, for demonstration see yellow bars in Fig 3C. The intensities of pixels (grey values) along this line were then determined by the “Profile plot tool” in FIJI, and profiles were scored as follows. The score “PM” (plasma membrane) was assigned to profiles that displayed two peaks in grey value near the cell periphery clearly distinguishable from the other signal (cytoplasmic signal), as shown by the representative plot in Fig 3C – WT. This distribution was typical for wild type LRT domain. The score “PM+” was assigned to profiles that displayed two peaks in grey value near the cell periphery and no or very little cytoplasmic signal, as shown in Fig 3C – L51F. This score meant greater plasma membrane association than the wild type. The score “PM-” was assigned to those profiles that showed two peaks in grey value near the cell periphery and a predominant signal between them. This score meant less plasma membrane association than the wild type (attenuated plasma membrane localization). Finally, the score “Cyto” (cytoplasmic) was assigned to those profiles that were characterized by a single broad peak in grey value, as shown by the representative plot in Fig 3C - L51N. This score meant no plasma membrane association. At least six cells from two independent experiments for each protein variant were analyzed to obtain a final score, which is shown in Figure 4 and Table 1.

### Hela cell fluorescence microscopy

HeLa cells at 40% confluence on coverslips were transfected with pEGFP-N2 (Clontech, USA) constructs encoding variants of BteA-GFP fusion protein (S3 Table) using Lipofectamine 2000 reagent (Invitrogen, USA). Eighteen h after transfection, HeLa cells were washed with PBS and fixed by 4 % formaldehyde solution in PBS (20 min, RT). The coverslips were then rinsed with distilled water and mounted onto a microscope glass slide using Vectashield mounting medium (Vector Laboratories, USA). Fluorescence microscopy was performed using an Olympus IX-81 inverted microscope with 60x UPlanSApo oil-immersion objective (N.A. =1.35). The camera and filter set were the same as for yeast cell imaging. Z-stacks were taken with 0.32 μm z-steps, and images collected in a 16-bit format were processed as during yeast cell imaging (deconvolution with Xcellence Advanced Maximum Likelihood (AMLE) filters and adjustment for brightness/contrast only). A single focal plane of a Z-stack is presented in all figures.

### Preparation of cell extracts and immunoblot analysis of GFP-tagged proteins

For preparation of yeast protein extracts, yeast cells with pYC2-CT vectors encoding BteA protein derivatives (S3 Table) were induced for 20 h by cultivation in SC-galactose media. Equivalents of OD_600_ = 1 of yeast cell cultures were collected, and denatured protein extracts were prepared by NaOH lysis/TCA precipitation method, according to [40].

Samples were mixed with SDS-PAGE sample loading buffer, heated for 5 min at 50 °C and separated on 10% SDS-PAGE gels. After the transfer onto nitrocellulose membrane, the proteins were probed overnight with rabbit anti-GFP antibody (1:2,000; clone D5.1, Cell Signaling Technology, USA) and revealed with horseradish peroxidase (HRP)-conjugated anti-rabbit IgG secondary antibody (1:3,000; GE Healthcare, USA). Blots were developed using a Pierce ECL chemiluminescence substrate (Thermo Fisher Scientific, USA) and an Image Quant LAS 4000 station (GE Healthcare, USA).

### Cytotoxicity assays

Cytotoxicity of *B. bronchiseptica* RB50 strain and its derivatives (S1 Table) towards HeLa cells was determined as a release of the intracellular enzyme lactate dehydrogenase (LDH) into the cell culture media using CytoTox 96 assay (cat.no. G1780, Promega, USA) or as changes in cell membrane integrity using CellTox Green Cytotoxicity Assay (cat.no. G8743, Promega, USA). Experiments were performed according to the manufacturer’s instructions. Briefly, 2 x 10^4^ HeLa cells per well were seeded into a 96-well plate in DMEM supplemented with 2% (vol/vol) FBS without the phenol red indicator and allowed to attach overnight. *B. bronchiseptica* cultures were grown to mid-exponential phase, and bacteria were added at the indicated multiplicity of infection (MOI). To enable efficient infection, plates were centrifugated (5 min, 400 g). For determination of LDH release, plates were incubated for 3 h at 37 °C and 5% CO_2_, which was followed by coupled enzymatic assay of cell culture media and absorbance measurement at 495 nm using Tecan Spark microplate reader (Tecan, Switzerland). The % of total LDH release was calculated using the following equation: (OD_495_ sample – OD_495_ media)/(OD_495_ total lysis-OD_495_ media)* 100. For measurement of cell membrane integrity, fluorescent DNA binding dye CellTox Green was added to HeLa cells in 96-well black clear bottom plate concomitantly with *B. bronchiseptica*. The plate was then placed inside the chamber with 37 °C and 5% CO2 of Tecan Spark microplate reader (Tecan, Switzerland), and fluorescent measurement at 490ex/525em was performed in 30 min intervals for 6 h.

### Molecular docking

Molecular docking was carried out using AutoDock Vina [41]. The structure of BteA-LRT (aa 29-121, PDB code: 6RGN) was processed by removing all sulfate ions and adding all hydrogens using AutoDockTools. The protein molecule was considered as a rigid molecule, whereas the ligand (inositol 4,5-bisphosphate) was treated as being flexible. The binding site region was defined as a region, which encompasses the entire surface of the protein. The binding energy was almost equal for both ligand binding sites, with the binding affinity value of −3.7 kcal/mol. The results of the docking were visualized using Chimera 1.14rc [42] and are shown in Fig 7B.

## Acknowledgments

We thank H. Lukeova, I. Marsikova, and L. Novakova for their excellent technical help. This work was supported by grant 18-16772Y of the Czech Science Foundation (www.gacr.cz) and project LM2018133 (Czech National Node to the European Infrastructure for Translational Medicine) of the Ministry of Education, Youth and Sports of the Czech Republic (https://www.msmt.cz) to J.K.

## Suppl. figure legends

**S1 Fig. Phospholipid binding by the N-terminal motif of *Bordetella* effector BteA.**

**(A)** Protein-lipid overlay assay. The recombinant GST-tagged N-terminal LRT domain (LRT) and full-length BteA (BteA/BtcA) protein of *B. pertussis* were incubated at 5 μg/ml with commercial lipid strips. The binding was detected using an anti-GST antibody followed by chemiluminescence detection. Recombinant GST was used as a control. Lysophosphatidic acid (LPA); lysophophocholine (LPC); phosphatidylinositol (PI); phosphatidylinositol phosphates (PIP, PIP2, PIP3); phosphatidylethanolamine (PE); phosphatidylcholine (PC); sphingosine-1-phosphate (S1P); phosphatidic acid (PA), and phosphatidylserine (PS).

**B)** SPR kinetic binding analyses of the interaction between GST and lipid vesicles. Serially diluted GST protein (at 500, 250, 125, 62.5, and 31.25 nM concentrations) was injected in parallel over the neutravidin sensor chip coated with the immobilized liposomes (100 nm in diameter) containing PC, PC/PS (80:20) or PC/PIP2 (95:5), and left to associate (120 s) and dissociate (380 s) at constant flow rate of 30 μl/min. For clarity, only the binding curve for the highest concentration of GST (500 nM) is shown.

**S2 Fig. Phospholipids PS and PIP2 guide plasma membrane association of *Bordetella* BteA effector and its LRT motif in cells.**

**(A-B)** Phospholipid levels in the yeast cells were monitored by localization of GFP-tagged lipid-specific probes. **(A)** PIP2-specific probe 2xPH(PLCδ) was used as a control of the specificity of decreased PS levels in the *cho1Δ* derivative of the *S. cerevisiae* BY4742, whereas **(B)** PS-specific probe GFP-Lact-C2 monitored the specificity of decreased PIP2 levels in the *mss4^ts^* of *S. cerevisiae* SEY6210 at the restrictive temperature. Representative images from two independent experiments with the same outcome are presented. Scale bar, 5 μm.

**(C)** GFP-tagged BteA (BteA-GFP) effector of *B. pertussis* was visualized upon galactose induction in the temperature-sensitive *mss4^ts^* mutant and wild type (WT) strain of *S. cerevisiae* SEY6210 after the shift from the permissive (25 °C) to restrictive temperature (38 °C). Representative images from two independent experiments with the same outcome are presented. Scale bar, 5 μm.

**S3 Fig. Leu51 residue is involved in hydrophobic interactions of the LRT motif with a phospholipid membrane.**

**(A)** Overlay plot of SPR sensograms obtained after injection of LRT, LRT-L51N, and LRT-L51F proteins at 250 nM concentration over the neutravidin sensor chip coated with the immobilized PC/PS (80:20) lipid vesicles. The binding curves are representative of five independent “one-shot kinetic” experiments.

**(B)** Western blot analysis. Protein extracts were prepared from yeast cell cultures expressing the indicated GFP-tagged LRT protein variants after 20 h induction with galactose. Equal volumes of extracts (0.4 ml of the culture equivalent; OD600 = 1) were separated on SDS-PAGE and analyzed by immunoblot using an anti-GFP antibody (1: 2,000). The arrow indicates the molecular weight of the intact LRT-GFP fusion protein.

**S4 Fig. Positively charged residues of the loop L1, helix B, and helix D are critical for the plasma membrane association of the LRT motif.**

*S. cerevisiae* BY4741 cells carrying plasmids encoding the indicated LRT-GFP protein variants were induced for 20 h for protein expression and examined by live-cell imaging. Representative images from two independent experiments with the same outcome are presented. Scale bar, 5 μm.

**S5 Fig. *Bordetella* BteA effector and its LRT domain exhibit a polarized localization in yeast cells.**

*S. cerevisiae* BY4741 harboring plasmids encoding GFP-tagged LRT domain (LRT-GFP) or full-length BteA (BteA-GFP) were cultivated for 20 h in the medium supplemented with galactose to induce protein expression followed by their live-cell imagining. Arrows in respective panels point to incipient bud sites, small buds and mother-bud necks of large buds. Scale bar, 5μm.

